# Extracellular Cation Binding Pocket Is Essential For Ion Conduction Of *Os*HKT2;2

**DOI:** 10.1101/471003

**Authors:** Janin Riedelsberger, Ariela Vergara-Jaque, Miguel Piñeros, Ingo Dreyer, Wendy González

**Author notes:** **Correspondence:** Janin Riedelsberger.

## Abstract

HKT channels mediate sodium uniport or sodium and potassium symport in plants. Monocotyledons express a higher number of HKT proteins than dicotyledons, and it is only within this clade of HKT channels that cation symport mechanisms are found. The prevailing ion composition in the extracellular medium affects the transport abilities of various HKT channels by changing their selectivity or ion transport rates. How this mutual effect is achieved at the molecular level is still unknown. Here, we built a homology model of the monocotyledonous *Os*HKT2;2, which shows sodium and potassium symport activity, and performed molecular dynamics simulations in the presence of sodium and potassium ions. By analyzing ion-protein interactions, we identified a cation binding pocket on the extracellular protein surface, which is formed by residues P71, D75, D501 and K504. Proline and the two aspartate residues coordinate cations, while K504 forms salt bridges with D75 and D501 and may be involved in the forwarding of cations towards the pore entrance. Functional validation via electrophysiological experiments confirmed the biological relevance of the predicted ion binding pocket and identified K504 as a central key residue. Mutation of the cation coordinating residues affected the functionality of HKT only slightly. Additional *in silico* mutants and simulations of K504 supported experimental results.

## 1 Introduction

A proper sodium (Na^+^)/potassium (K^+^) homeostasis is a crucial requirement for high yielding plant growth. Although Na^+^ ions can promote plant growth at low concentrations, they turn hazardous at increasing levels (Leigh and Wyn Jones, 1984; Maathuis, 2009; Marschner, 2012). Due to similar physicochemical properties, Na^+^ ions can mimic functions of K^+^ ions, bridging periods of K^+^ shortage (Benito et al., 2014; Horie et al., 2007). However, at high concentrations, Na^+^ ions compete with K^+^ and cause K^+^ deficiency symptoms in plants, since Na^+^ imitates K^+^’s functions only incompletely (Hasegawa et al., 2000; Pardo and Quintero, 2002).

Plants have developed sophisticated mechanisms to cope with salt stress. These systems aim to avoid high cytosolic Na^+^ concentrations in cells of plant shoots by compartmentalization and retrieval of Na^+^ from the xylem sap (Adams and Shin, 2014; Flowers and Läuchli, 1983). HKT channels form one comprehensive family that is involved in Na^+^ usage and detoxification (Horie et al., 2009; Maser et al., 2002; Munns et al., 2012; Møller et al., 2009). In monocots, like rice, two types of HKT channels exist: (i) class I-type channels, which are mainly Na^+^ selective, and (ii) channels of class II acting as Na^+^/K^+^ co-transporters (Almeida et al., 2013; Garciadeblás et al., 2003). Among HKTs, there is a wide variety of transport kinetics, cation selectivity and rectification properties even within the two classes (Gassmann et al., 1996; Haro et al., 2005; Horie et al., 2001; Jabnoune et al., 2009; Rubio et al., 1995; Yao et al., 2010).

Several studies have demonstrated the interactive effect of different ion species for various HKT channels. In particular, the mutual effects of Na^+^ and K^+^ have been studied (Horie et al., 2001; Oomen et al., 2012; Yao et al., 2010). However, the mechanism by which ion species affect each other and modulate the ion transport process is so far not understood. Initially, it was discussed that HKT channels form multi-ion pores (Gassmann et al., 1996; Yao et al., 2010). In this scenario, multiple ion binding sites exist in the pore allowing a coupled ion transport without additional conformational changes (Su et al., 1996). However, recently Böhm and colleagues experimentally demonstrated that in the Venus flytrap HKT1 a maximum of one ion at a time occupies the selectivity filter (Böhm et al., 2015). Therefore, the mechanism behind mutual ion species effects remains controversial.

In this study, we provide new functional-structural perspectives for the understanding of ion species-specific effects on ion conduction in HKT channels. We built a homology model of the *Oryza sativa* HKT channel *Os*HKT2;2 embedded in a lipid membrane, performed molecular dynamics simulations in the presence of Na^+^ and K^+^ ions and evaluated protein-ion interactions. Through systematic analysis, we identified a potential cation binding pocket at the extracellular surface of the protein – a region that ions have to pass before entering the pore. The cation binding pocket was characterized by computational studies and its functional importance for *Os*HKT2;2 was experimentally validated using electrophysiological methods. Additionally, analyses of *in silico* mutants further underpinned the experimental insights.

## 2 Materials and Methods

### 2.1 Homology modelling and molecular dynamics simulation

The structural model of *Os*HKT2;2 was generated using the I-Tasser server (Roy et al., 2010). Full-length structural models were built from multiple threading alignments. For threading the bacterial channels KtrB (PDB ID 4J7C) and TrkH (PDB ID 3PJZ) were used. The N-terminus of *Os*HKT2;2 (residues 1-38) was modeled on the basis of TrkH, since the crystal structure of KtrB does not contain the N-terminal region. *Os*HKT2;2 contains a 29 amino acid long extracellular loop that is not present in bacterial channels. This loop spanning residues 473-501 has been modeled separately using the FALC-Loop server, which combines statistical and knowledge-based methods (Ko et al., 2011). The improved full-length model was refined using i3Drefine (Bhattacharya and Cheng, 2013) and minimized using CHARMM via its web portal CHARMMing (Brooks et al., 2009; Miller et al., 2008).

The refined model was embedded in a pre-equilibrated POPC bilayer solvated with preequilibrated TIP3P water molecules (Jorgensen et al., 1983) in a periodic boundary condition box using VMD (Humphrey et al., 1996). The system was neutralized by the addition of sixteen chloride ions. Each, six Na^+^ and K^+^ ions as well as twelve further chloride ions were added to mimic the concentration of 10mM NaCl and 10mM KCl. The entire system underwent several minimization steps and was equilibrated with successive restraint reduction until restraints were removed entirely. Three replicates of 100ns molecular dynamics (MD) simulations were generated using NAMD (Kalé et al., 1999; Phillips et al., 2005).

Binding pocket residues were identified by counting the frames in which an ion was within 4Å of a given residue during MD simulations. Ion coordination was examined by measuring distances between charged atoms of P71, D75, D501, K504 and the ion present in the binding pocket. Salt bridges were evaluated using VMD’s Salt Bridge plugin. Electrostatic potentials were calculated using the APBS web server with the ionic strength set to simulate 10mM NaCl and 10mM KCl (Unni et al., 2011). APBS input files were generated using the PDB2PQR Server version 2.1.1 (Dolinsky et al., 2004; 2007).

*In silico* mutations of *Os*HKT2;2 were generated via VMD’s Mutate plugin. Mutants were minimized and equilibrated followed by 15ns of MD simulation in NAMD.

### 2.2 Electrophysiology

Wild-type and mutant *Os*HKT2;2 constructs were cloned into the pNB1u vector for expression in *Xenopus* oocytes using the USER-cloning technique (Nour-Eldin et al., 2006). Utilized primers were: GGCTTAAUatgacgagcatttaccaagaa (forward) and GGTTTAAUctaccatagcctccaatatt (reverse). Vector-specific sequences are given in uppercase format, *Os*HKT2;2-specific sequences in lowercase format. The position of uracil is underlined. Standard fusion PCR technique was used for site-directed mutagenesis. After DNA linearization with *NotI*, cRNA was synthesised using the mMessage mMachine *in vitro* transcription kit following the manufacturer’s guidelines.

For expression in oocytes, stage V and VI *Xenopus leavis* oocytes were harvested and kept in ND96 solution (96mM NaCl, 2mM KCl, 1.8mM CaCl2, 1mM MgCl2, 2.5mM Na Pyruvate, 5mM Hepes – pH 7.5, 50 mg mL^−1^ gentamycin and 0.4g L^−1^ BSA). Oocytes were defolliculated by collagenase treatment in ND96 without CaCl_2_, gentamycin and BSA and kept overnight at 18°C in complete ND96 solution as described initially. All animal procedures were performed in accordance with Cornell University IACUC Protocol number 2017-0139. 50nl of cRNA (500ng/μl) were microinjected into oocytes using an oil-driven injection system (Nanoject II Auto-Nanoliter Injector, Drummond Scientific Company, US). Cells were then incubated for two days at 18°C in complete ND96 solution. Whole-cell currents were recorded using conventional Two-Electrode Voltage-Clamp technique (GeneClamp 500 amplifier and Digidata 1320A-PClamp 10 data acquisition system, Axon Instruments). Recordings were carried out under constant perfusion of bath solutions containing 2mM MgCl_2_, 1.8mM CaCl_2_, 10mM MES, pH 5.5 with Tris-Base with the addition of: Na0K1: 1mM KCl, 165mM Sorbitol; Na0K30: 30mM KCl, 130mM Sorbitol; Na03K0: 0.3mM NaCl, 165mM Sorbitol; Na03K1: 0.3mM NaCl, 1mM KCl, 165mM Sorbitol; Na30K0: 30mM NaCl, 130mM Sorbitol; Na30K1: 30mM NaCl, 1mM KCl, 130mM Sorbitol; Na30K30: 30mM NaCl, 30mM KCl, 100mM Sorbitol.

## 3 Results

### 3.1 Electrophysiological characterization of *Os*HKT2;2

*Os*HKT2;2 is one out of eight to nine HKT channels (dependent on the cultivar) expressed in rice plants (Garciadeblás et al., 2003). Members of the HKT family show remarkable functional diversity regarding ion selectivity, rectification properties and effect of external cation compositions (Jabnoune et al., 2009). To provide a solid basis for this study, we first characterized the functional properties of *Os*HKT2;2 expressed in *Xenopus leavis* oocytes using the Two-Electrode Voltage-Clamp (TEVC) technique.

Under voltage clamp conditions, cells expressing *Os*HKT2;2 conducted mainly Na^+^ inward currents in the absence of K^+^. Thereby, ion transport rates raised with increasing Na^+^ concentrations (Figure 1B, D, I). At low extracellular Na^+^ concentrations (0.3mM NaCl), ion currents of several hundred nanoamperes were detected, while at high Na^+^ concentrations (30mM NaCl) currents increased to several microamperes consistent with Na^+^ being the ion carrying the current. A 100-fold increment of Na^+^ concentration (from 0.3mM to 30mM) resulted in an almost 5-fold increase of ion conduction.

**Figure 1:**
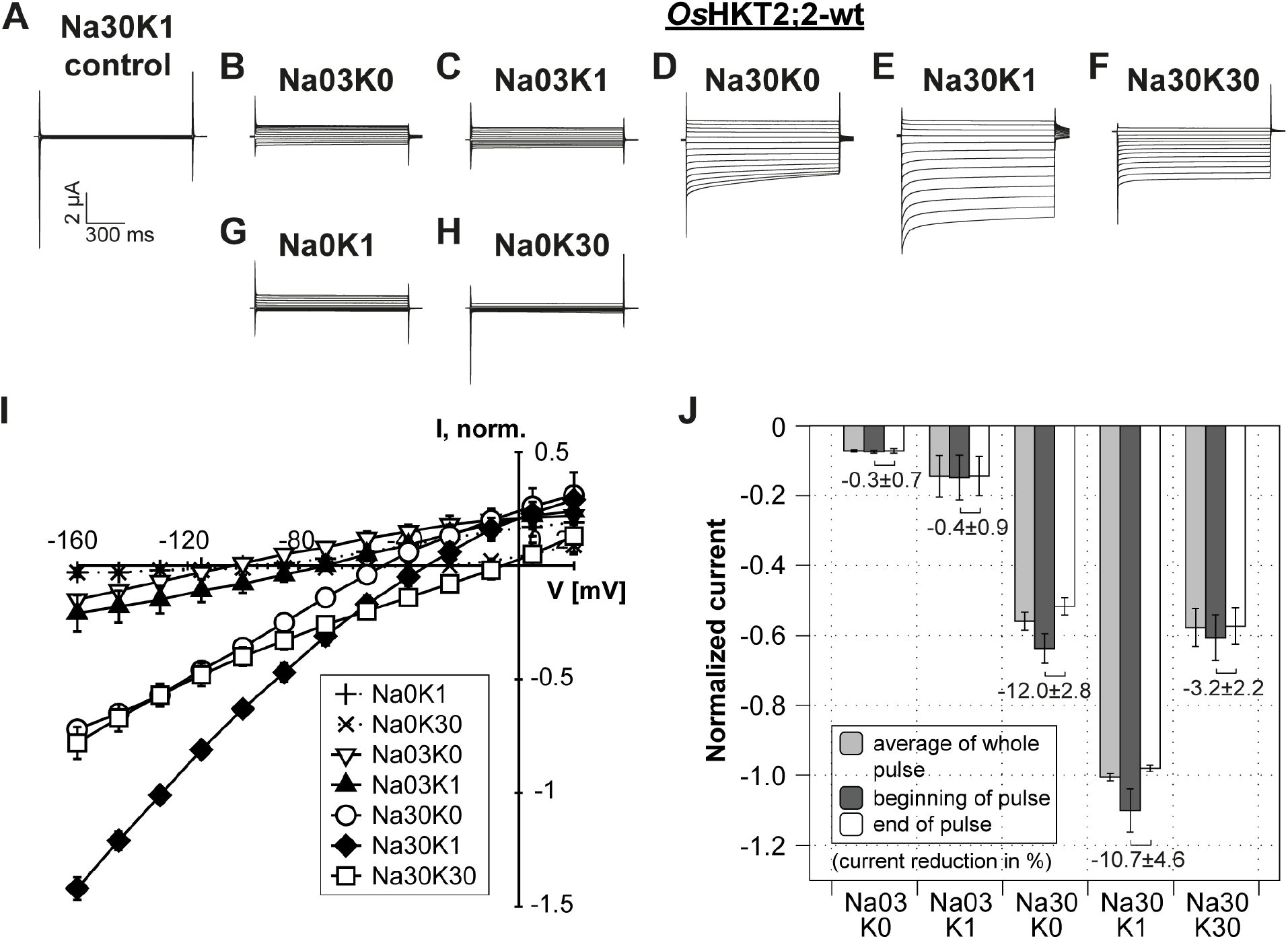
Electrophysiological characteristics of *Os*HKT2;2 in *Xenopus* oocytes. (**A**) Representative currents elicited in control cells (not injected with cRNA). Only currents for Na30K1 solution are shown since highest currents are expected under these conditions. Control currents for all conditions are shown in Supporting Figure 3. (**B-H**) Representative *Os*HKT2;2 mediated whole cell currents recorded two days after cRNA injection. Currents were recorded at indicated Na^+^ and K^+^ concentrations: Na0K1 – no NaCl and 1mM KCl, Na0K30 – no NaCl and 30mM KCl, Na03K0 - 0.3mM NaCl without KCl, Na03K1 - 0.3mM NaCl and 1mM KCl, Na30K0 - 30mM NaCl without KCl, Na30K1 - 30mM NaCl and 1mM KCl, Na30K30 - 30mM NaCl and 30mM KCl. The holding potential was set to the zero current level (approximately - 45mV with solution Na30K1) and 1s voltage pulses were stepped from +20 to −160mV in −15mV decrements. A 1.5s resting interval was allowed between successive voltage steps. The current (μA) and time (ms) scale for all traces shown in A through H is shown at the bottom of panel A. (**I**) Mean current-voltage (IV) curves. Standard deviation calculated from data obtained from five to eight measurements obtained on five different days and from four different oocyte batches. Currents were normalized to currents measured at −145mV in Na30K1 solution. (**J**) Currents elicited by a −145mV voltage pulse, measured at the beginning (dark grey) or end (white) of the voltage pulse, and as average currents over the whole voltage pulse (light grey). Currents were normalized to currents measured in Na30K1 solution. Numbers in the plot indicate the degree of current reduction over time (from beginning to end of the voltage pulse). The magnitude of deactivation is stated in percentage. Means and standard deviation were obtained from data of five to eight measurements.

The addition of low K^+^ concentrations (1mM KCl) had a stimulating effect on the ion transport and led to a significant increase in ion conduction (Figure 1D, E). One millimolar of KCl almost doubled the ion conduction at high and low Na^+^ concentrations. We are assuming a stimulating effect of K^+^ ions for two reasons: (1) In the absence of NaCl in the bath solution, no inward currents could be measured in the presence of 1 or 30mM KCl (Na0K1, Na0K30). Currents at negative voltages were comparable to measurements in control oocytes (compare Figure 1G, H, I and Supplemental Figure 3). Therefore, potassium ions, at least in the absence of sodium ions, do not pass the conduction pathway. Hence, it is unlikely that the presence of 1mM KCl is simply adding to the current of 30mM NaCl. (2) This stimulating effect was revoked under rising K^+^ concentration (30mM KCl; Figure 1F, I, J). Interestingly, the ionic current at 30mM KCl was comparable to that observed without K^+^ ions in the extracellular solution indicating that K^+^ stimulates ion conduction only at low concentrations.

Additionally, under high Na^+^ concentrations, inward currents partially deactivated at negative voltages. This effect was most pronounced in the absence of K^+^ and reduced with increasing K^+^ concentrations (Figure 1D-F, J). At 30mM NaCl (no KCl), the current amplitude was reduced by 12.0% (±2.8%) over one-second voltage pulses. In the presence of 1mM KCl, the current reduction was 10.7% (±4.6%), and at 30mM KCl current deactivation was almost abolished (3.2%±2.2%).

Overall, we identified and characterized three functional properties for wild-type *Os*HKT2;2 which we used later in the study to compare functional changes in mutated *Os*HKT2;2 channels (Table 1). The functional properties of wild-type *Os*HKT2;2 include: (1) ionic currents enhance with increasing Na^+^ concentration, (2) low K^+^ concentration (1mM KCl) stimulates Na^+^ ion transport, and (3) high K^+^ concentration (30mM KCl) lacks the stimulating effect on Na^+^ ion transport.

**Table 1:**
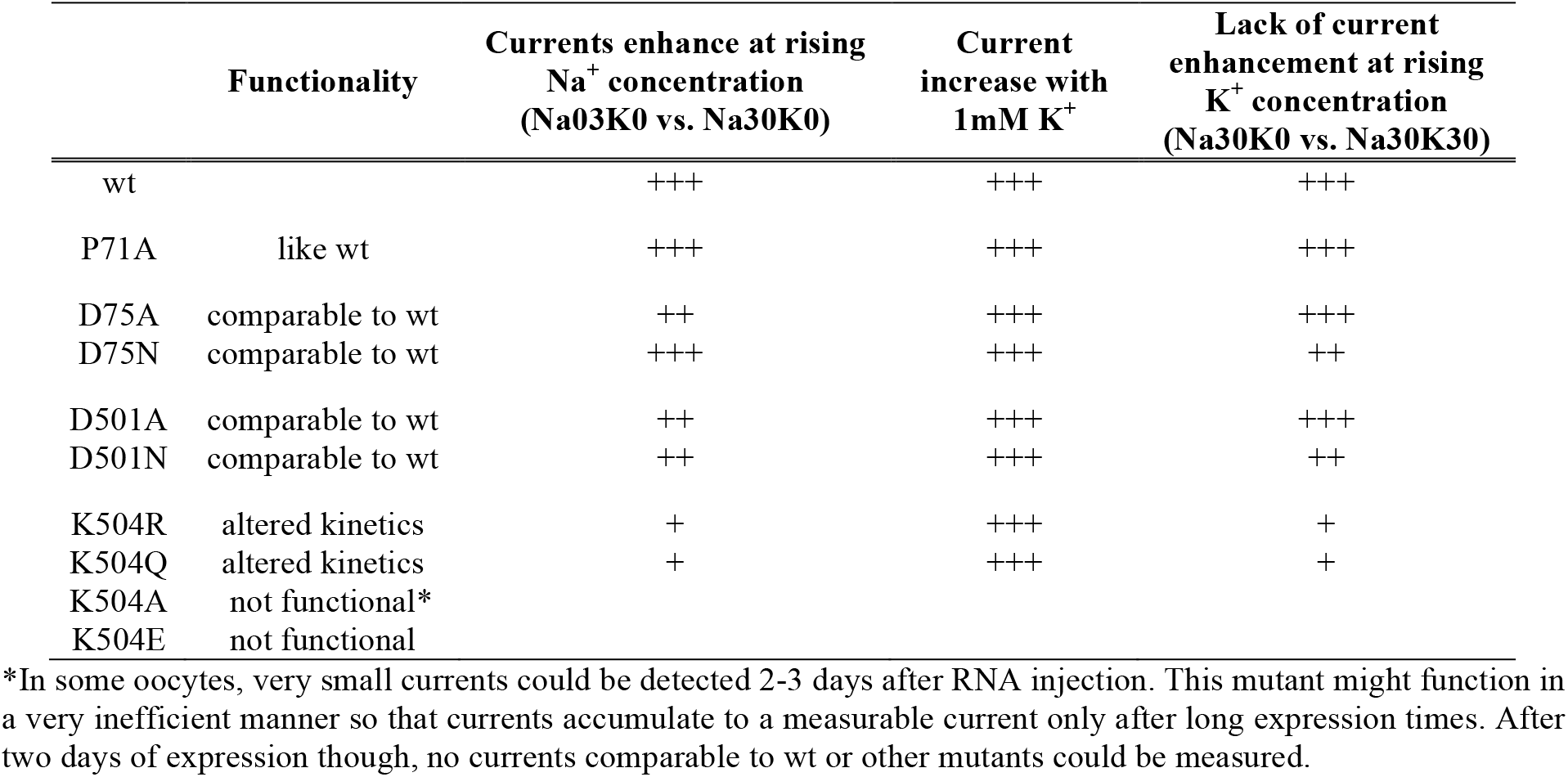
Electrophysiological characterisation of *Os*HKT2;2 and its mutants. The presence of three current characteristics are specified according to their degree of occurrence: very strong (+++), medium (++) or weak (+).

### 3.2 Identification of an extracellular cation binding pocket

In order to identify amino acids that are approached by ions before entering the pore, we built a homology model of the *Os*HKT2;2 wild-type channel on the basis of the structurally and functionally related bacterial channels KtrB (PDB ID 4J7C) and TrkH (PDB ID 3PJZ). Because the N-terminus of KtrB is not included in the crystal structure, N-terminal amino acids of *Os*HKT2;2 were modeled on the basis of the TrkH (see Methods for details). Subsequently, we performed molecular dynamics (MD) simulation in the presence of 10mM NaCl and 10mM KCl for 100ns. This allows the evaluation of the frequency at which ions approach residues over time. To demonstrate reproducibility, the MD simulation was performed threefold and a total of 300ns simulation time was reached, which form the basis of the following results.

Analyses of the contact frequency between ions and amino acids during MD simulations led to the identification of a potential extracellular cation binding pocket. This binding pocket is located in the outer extracellular protein region approximately 20Å away from the pore entrance (Figure 2A). We identified three polar residues and one proline to be involved in the formation of a putative cation binding pocket – P71, D75, D501 and K504. Within these, three are highly conserved among plant HKT channels (P71, D75 and K504), while at position of D501 a negative charge is preserved in 75% of HKT channel sequences. Figure 2B illustrates the conservation of these four residues within the rice HKT family and additionally states their conservation within a multiple sequence alignment based on 20 experimentally characterized HKT channels.

**Figure 2:**
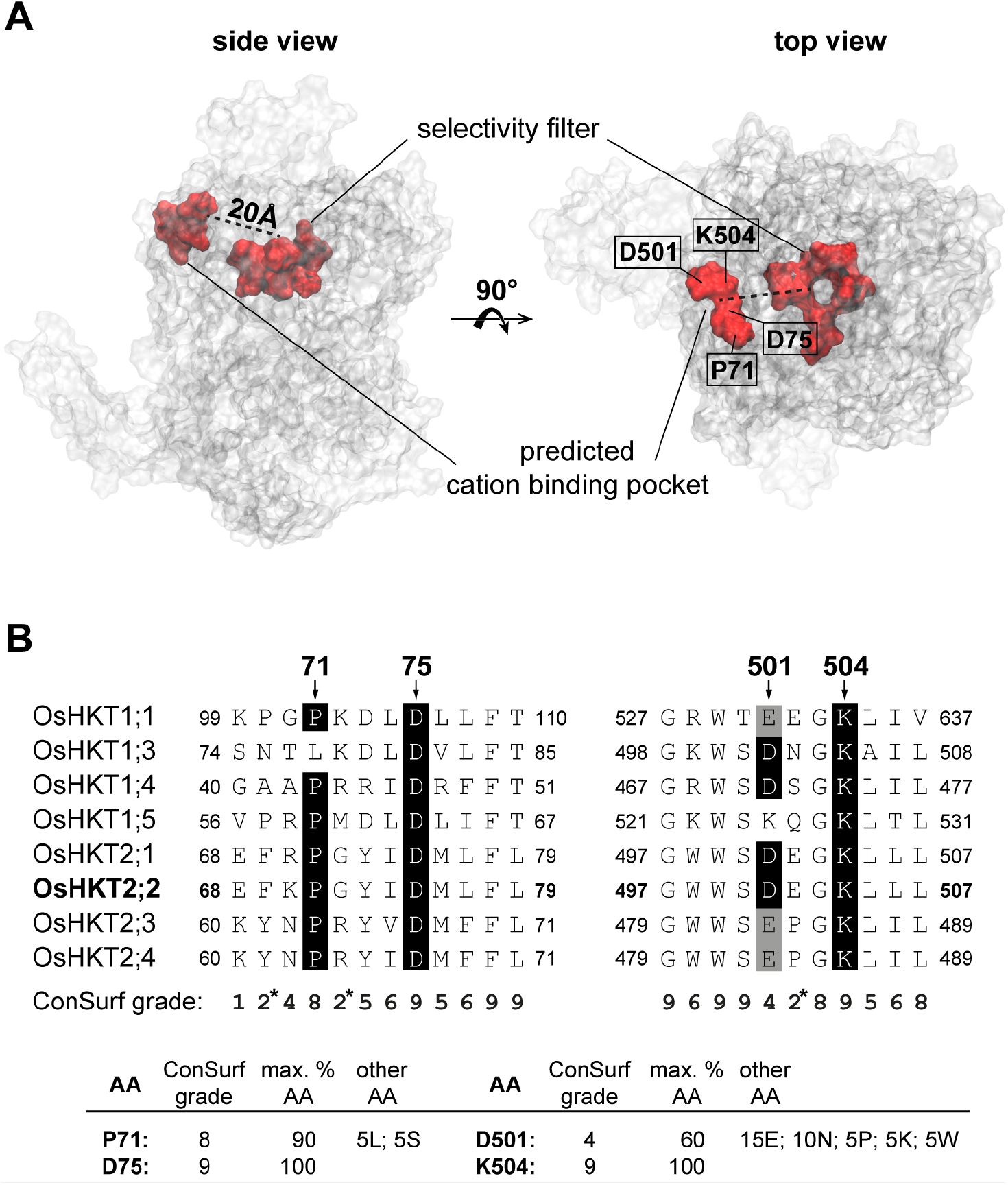
Localization and conservation of extracellular cation binding pocket. (**A**) *Os*HKT2;2 is displayed as transparent surface representation with the selectivity filter and surrounding residues marked in the center of the channel as red surface representation. The cation binding pocket is marked on the outer extracellular edge of the protein also as red surface representation, and the position of the four contributing residues is shown. The approximate distance between binding pocket and pore is indicated in the top view representation. (**B**) An extract of the multiple sequence alignment of all eight HKT members present in *Oryza sativa*. Conservation of the residues P71, D75, D501 and K504 forming the binding pocket is indicated. ConSurf grades were calculated on the basis of a multiple sequence alignment of 20 experimentally characterized HKT sequences, and are shown below the sequence alignment. The grades range from 1 (variable) to 9 (conserved) (Landau et al., 2005). D75 and K504 are 100% conserved (ConSurf grade 9). P71 is in 90% of HKT sequences conserved (ConSurf grade 8), and contains in 5% of the sequences a lysine or serine. At position 501 a negative charge is somewhat conserved as aspartic in 60% of the sequences (ConSurf grade 4) or glutamic acid (in 15% of the sequences). In the remaining 25% of the sequences, this position holds an asparagine (10%), proline (5%), lysine (5%), or tryptophan (5%).

#### 3.2.1 Structural attributes of the extracellular cation binding pocket

Both, sodium and potassium ions, approached a defined area on the protein surface during MD simulations. K^+^ ions were found in this region between 0.5 to 1% of the simulation time, while Na^+^ stayed there about three times longer (1.2 to 5.2% of the time; Figure 3A). At this point, it is important to clarify that we were not searching for a region in the sense of an ion binding site that binds ions for extended time periods with potential allosteric impacts on protein function. Instead, we were searching for areas, which ions approach and pass on their way into the pore – in other words, binding sites that attract cations and guide them to the pore entrance. That implicates that the total time of residence of an ion within the identified region is comparatively short as observed in our MD simulations. Nevertheless, cations approach the site frequently and are coordinated by the identified residues, in contrast, to randomly approached residues where no ion coordination occurs. Throughout the remaining of the study, we will refer to this region as extracellular cation binding pocket.

**Figure 3:**
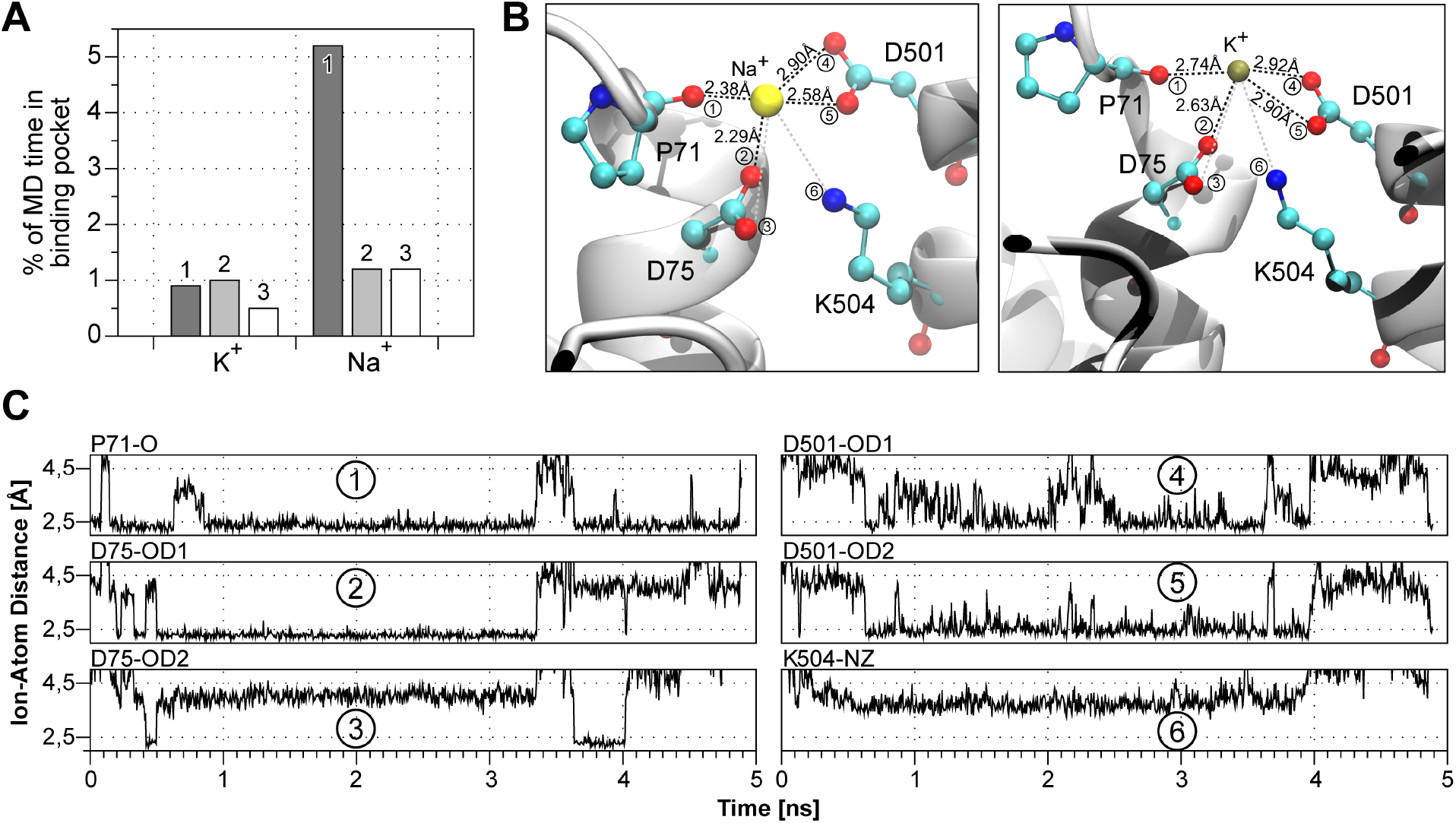
Oxygen atoms of P71, D75 and D501 coordinate cations. (**A**) Total time that K^+^ and Na^+^ stay in the binding pocket during each of the three simulations expressed in percentage. (**B**) Coordination of cations happens via the backbone oxygen atom of P71, one side chain oxygen atom of D75 and both side chain oxygen atoms of D501 (red spheres). Average ion-atom distances for sodium and potassium are indicated in the figure. Sodium: 1) 2.38±0.14Å, 2) 2.29±0.09Å, 3) 4.02±0.21Å, 4) 2.90±0.65Å, 5) 2.58±0.31Å and 6) 3.74±0.25Å. Potassium: 1) 2.74±0.19Å, 2) 2.63±0.30Å, 3) 4.15±0.40Å, 4) 2.92±0.47Å, 5) 2.90±0.38Å and 6) 3.96±0.28Å. (C) Exemplary plot of ion-atom distances for sodium. Distances of one sodium ion and all six charged atoms facing the potential binding pocket are represented for 5ns of MD system 1 while Na^+^ is inside the pocket. The close proximity (<2.5Å) of Na^+^ to the backbone oxygen of P71, one side chain oxygen of D75 and both side chain oxygen atoms of D501 is illustrated, indicating their ability to coordinate sodium.

Despite their short residence in the binding pocket, the cations are coordinated by surrounding amino acids. A closer examination revealed four oxygen atoms coordinating the cations in this region: the backbone oxygen of P71 (Figure 3B, C.1), one side chain oxygen atom of D75 (Figure 3B, C.2) and the two side chain oxygen atoms of D501 (Figure 3B, C.4 and C.5). The average distances between the mentioned oxygen atoms and Na^+^ were 2.38Å, 2.29Å, 2.90Å and 2.58Å, respectively, and, 2.74Å, 2.63Å, 2.92Å and 2.90Å for K^+^ (Figure 3B). The coordination of one Na^+^ ion over time is exemplarily shown in Figure 3C. Besides the proximity (<2.5Å) of the Na^+^ ion to the four mentioned oxygen atoms (plots C.1, C.2, C.4, C.5), the proximity to other side chain atoms facing towards the predicted binding pocket is also shown. Namely, the proximity to the second side chain oxygen atom of D75 (plot C.3) and the positively charged nitrogen atom of K504 (plot C.6). The second side chain oxygen atom of D75 is oriented away from the binding pocket most of the time and cannot contribute to the coordination of cations. As expected, the nitrogen of lysine 504 is also not involved in the coordination, since the positive charges of cations and the nitrogen atom repel each other.

Nevertheless, lysine 504 is located between the two aspartic acids D75 and D501, which suggests an interaction between these residues. Salt bridges may be formed between oppositely charged residues when the charged atoms are closer than 4Å (Kumar and Nussinov, 2002). Average distances between the positively charged nitrogen atom of lysine 504 and the negatively charged oxygen atoms of both aspartates fulfill this requirement (Figure 4A, B). Indeed, both oxygen atoms of D75 are in constant proximity to the nitrogen of lysine 504 (Figure 4B, upper two traces) with an average distance of 2.83Å and 2.82Å, respectively (Figure 4C). The two oxygen atoms of D501 alternate in their proximity to this nitrogen (Figure 4B, lower two traces). If in proximity, the distance is 2.68Å on average (Figure 4C). Thus, lysine 504 may interact with both aspartates and acts as a structural component to obtain the geometry of the binding pocket by keeping the aspartates in place rather than coordinating ions.

**Figure 4:**
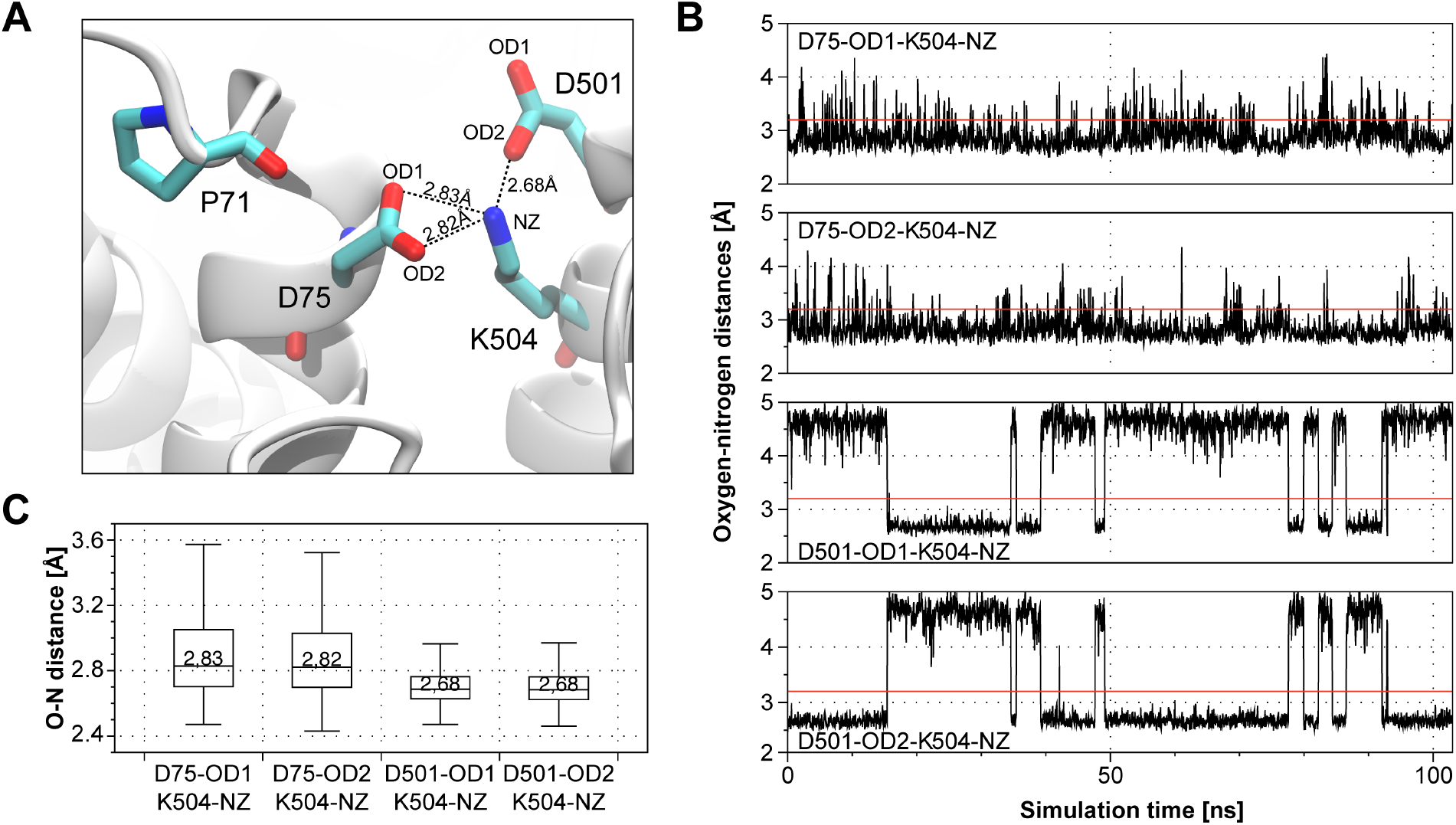
K504 can form salt bridges with D75 and D501. (**A**) Representation of potential salt bridges between K504 and D75 and D501. The median oxygen-nitrogen distance is stated. (**B**) Exemplary plot of oxygen-nitrogen distances throughout one complete simulation. The formation of salt bridges is considered possible when oxygen-nitrogen distances are below the threshold of 3.2Å. The red line in each plot indicates this threshold. (**C**) Distance distribution of indicated oxygen-nitrogen distances over all three simulations. The median is stated in each box plot. Outliers are not drawn. For each of the two D501 oxygen atoms only those time spans where the respective oxygen atom was orientated towards K504’s nitrogen atom were used.

Overall, our computational studies suggest that the extracellular cation binding pocket is constituted by a positively charged lysine (K504) that holds two negatively charged aspartates (D75 and D501) in place, which form a negative environment that attracts cations. The side chain oxygen atoms of both aspartates, in conjunction with the backbone oxygen of P71 coordinate Na^+^ and K^+^ ions.

### 3.3 Experimental validation of the predicted cation binding pocket

To study the relevance of the cation binding pocket for the ion conduction of *Os*HKT2;2, we mutated P71, D75, D501 and K504 and experimentally characterized the changes in functionality in these mutants. Alanine mutations of all four residues were performed to assess the effect of side chain removal, and asparagine mutants of D75 and D501 allowed the evaluation of negative charges neutralization. Besides, K504 was mutated to an arginine to examine the relevance of the side chain size, to glutamine to analyze the effect of charge neutralization, and to glutamic acid to evaluate the inversion from a positive to a negative charge. Overall, mutation of residue K504 had the most significant impact on *Os*HKT2;2’s functionality. Modifications of D75 and D501 affected the kinetic features slightly, whereas mutation of P71 did not cause changes in the transport mechanism as compared to the wild-type channel (Table 1).

#### 3.3.1 Mutations of P71, D75 and D501 show no or minor functional alterations

To evaluate the effect of mutations of the cation coordinating residues we modified P71, D75 and D501 by the removal of side chains (alanine mutants) and charges (asparagine mutants). The mutant P71A behaved similarly to the wild-type channel regarding the three functional properties that have been established before on the basis of wild-type *Os*HKT2;2: Na^+^ conductivity (1), the mutual ion-species effect at low (2) and high (3) K^+^ concentrations (Table 1). P71A transported Na^+^ in the absence of K^+^, and the ion transport increased at high Na^+^ concentration. As observed in the wild-type channel, ion currents increased in the presence of 1mM K^+^ but not 30mM K^+^, when compared to currents recorded in the absence of K^+^ (Figure 5).

**Figure 5:**
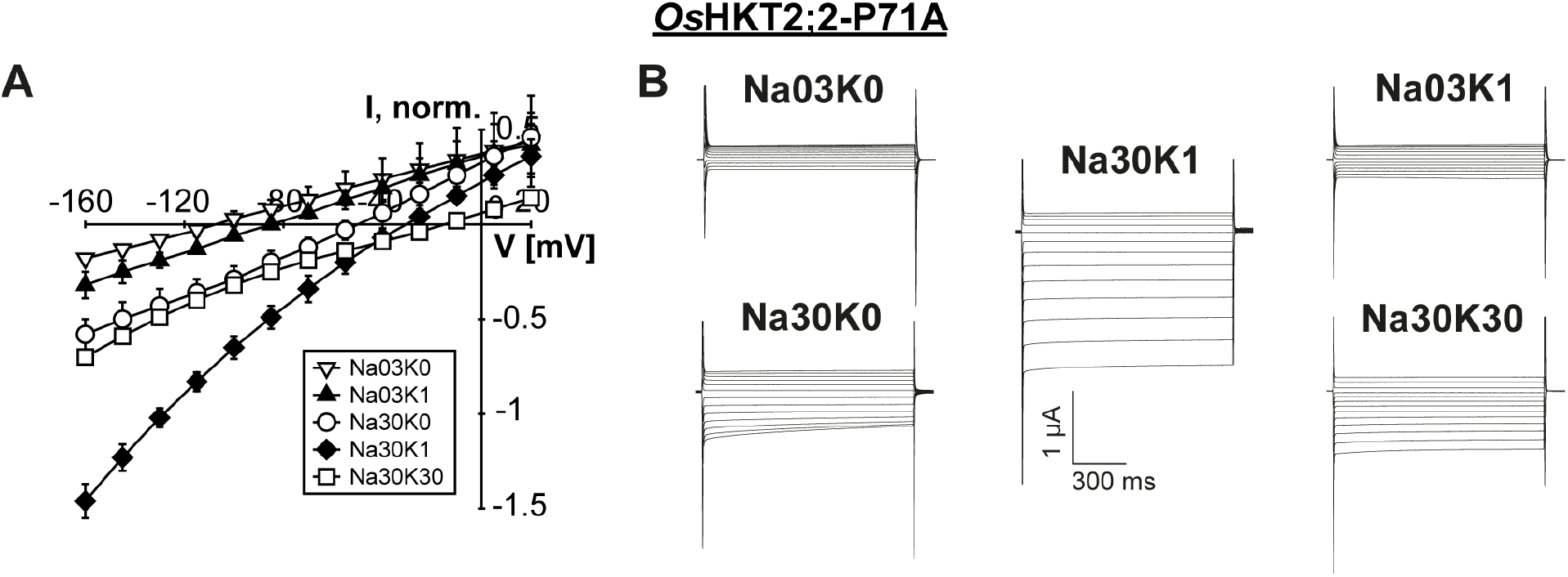
Mutation of P71 does not affect channel functionality. (**A**) Mean current-voltage (IV) curves and standard deviation calculated from data obtained from at least three different oocytes. (**B**) Representative currents measured at the indicated NaCl and KCl concentrations in *Xenopus* oocytes two days after cRNA injection. A pulse at holding potential (zero current level) was followed by 1s voltage pulses from +20 to −160mV in −15mV decrements and continued with a final pulse at holding potential for 1.5s.

Mutations of residues D75 and D501 affected the ion conduction slightly. As observed in the wild-type channel, D75A/N and D501A/N transported Na^+^ in the absence of K^+^ and a concentration-dependent manner. However, the increase of the current amplitude was slightly lower for D75A and D501A (4-fold), and only 3-fold in the D501N mutant (Figure 6). The strong stimulating effect of low K^+^ concentrations on the ion conduction was observed in all mutants. However, in contrast to wild-type *Os*HKT2;2, low magnitudes of current activation at 30mM NaCl were still observed in D75N and D501N mutants (Figure 6C, F, G and Supporting Figure 1). Indicating that the mutual effect of K^+^ on Na^+^ currents had slightly shifted.

**Figure 6:**
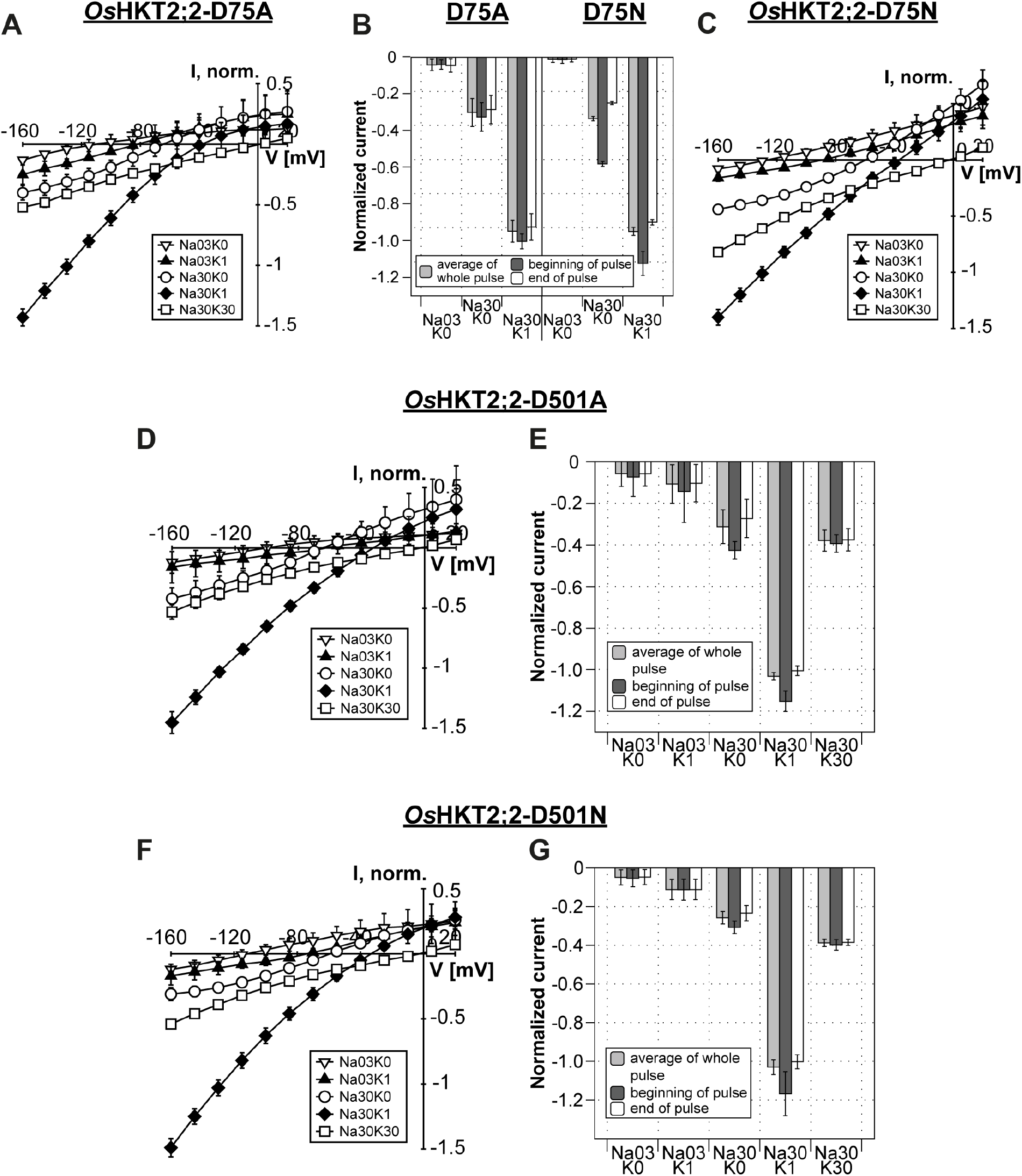
Mutations of D75 and D501 maintain channel functionality with only small changes in the kinetics. (**A, C, D, F**) Mean current-voltage (IV) curves and standard deviation calculated from data obtained from at least three different oocytes. (**B, E, G**) Currents measured at −145mV at the beginning (dark grey) or end (white) of voltage pulses and of average currents over the whole voltage pulse (light grey) of D75A and D75N (**B**), D501A (**E**) and D501N (**G**). Currents were normalized to currents measured in Na30K1 solution. Means and standard deviations were obtained from data of three to five different oocytes.

In summary, residues D75 and D501 were more sensitive towards modifications than P71, which was not affected by mutation. The latter was expected since proline coordinates cations via its backbone oxygen atom, which is not altered by mutations.

#### 3.3.2 K504 mutants showed substantial differences in their functionality compared to wild-type *Os*HKT2;2

Lysine 504 was identified as a structural component of the cation binding pocket that seems to take a crucial position in the overall geometry. Indeed, K504 was very sensitive towards residue substitutions. Removal of the side chain (mutant K504A) and inversion of the charge (mutant K504E) rendered *Os*HKT2;2 non-functional. K504A and K504E injected oocytes showed no currents two days after cRNA injection (Figure 7E, F). Each, fifteen injected oocytes have been analyzed on three different measuring days. However, oocytes injected with K504A cRNA sporadically conducted low currents in the nanoampere range, which differed significantly from those recorded in control oocytes. These observations were occasionally made three days after cRNA injection. We speculated that the K504A mutant may be functional but conducts ions with extremely low efficiency. In this case, the increased protein expression over time may lead to increased whole-cell currents through *Os*HKT2;2-K504A that are only detectable after long incubation times.

**Figure 7:**
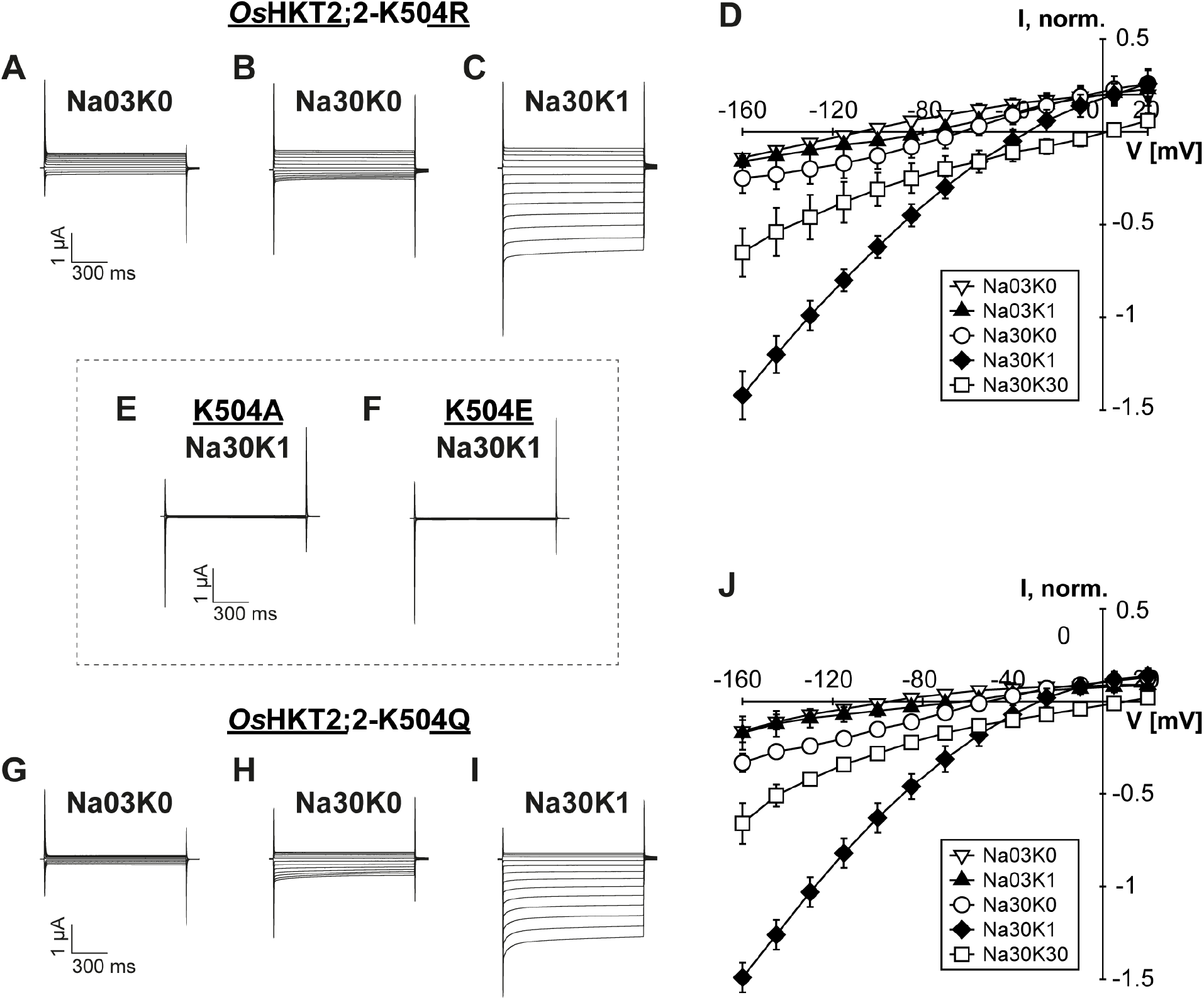
Mutations of K504 affect conduction behaviour of Na^+^ substantially. (**A-C, E-I**) Representative currents recorded in *Xenopus* oocytes two days after cRNA injection at indicated Na^+^ and K^+^ concentrations: Na03K0 - 0.3mM NaCl without KCl, Na30K0 - 30mM NaCl without KCl, Na30K1 - 30mM NaCl and 1mM KCl. A pulse at holding potential (zero current level) was followed by 1s voltage pulses from +20 to −160mV in −15mV decrements and continued with a final pulse at holding potential for 1.5s. (**D, J**) Mean current-voltage (IV) curves and standard deviation calculated from data obtained from three to six measurements.

In contrast, lysine mutants K504R and K504Q were functional although with substantial differences in their transport characteristics as compared to wild-type *Os*HKT2;2 and all other mutants. Na^+^ currents in the absence of K^+^ were low even at high Na^+^ concentrations (30mM NaCl). In fact, the current amplitude rose not more than 2-fold with increasing Na^+^ concentration (compared to the 5-fold increase in wild-type channels) and remained in the nanoampere range. Therefore, the increase of external Na^+^ concentration cannot enhance the ion conduction in the same extent as in wild-type *Os*HKT2;2 (Figure 7B, D, H, J). Nevertheless, low K^+^ concentrations did maintain their strong stimulating effect on the ion conduction (Figure 7C, I). This effect was even more pronounced in the lysine mutants. In the background of 30mM NaCl, the addition of 1mM KCl resulted in a more than 4-(K504Q) to 6-fold (K504R) increase of the current amplitude. Furthermore, the increase of K^+^ concentration in the extracellular bath media had still a substantial stimulating effect on the ionic conduction. Currents measured at 30mM Na^+^ and K^+^ were about twice as large in amplitude compared to currents in the presence of 30mM Na^+^ only (Figure 7D, J). This observation indicates that the Na^+^ transportability of *Os*HKT2;2 is affected in the K504R and K504Q mutants, while K^+^ ions are still able to promote and, in these mutants, also to rescue proper ion conduction.

### 3.4 *In silico* mutations support experimental results

To gain further understanding of the nature of the experimental results, the four lysine mutants were studied *in silico* by performing 100ns MD simulations. In each simulation, we examined the time Na^+^ and K^+^ ions approached one of the four binding pocket residues (<4Å) and the total number of approaches throughout the MD simulation (Table 2). The combined assessment of duration of stay and number of approaches allows the evaluation of the average time an ion spent in the cation binding pocket. It is striking that Na^+^ ions approached the binding pocket of the non-functional lysine mutants K504A and K504E more frequently than in the functional mutants K504Q and K504R (35 and 85 times versus 15 and 16 times; Table 2). The approaches observed in the mutants K504Q/R were comparable to the number of Na^+^ ion approaches in wild-type *Os*HKT2;2.

**Table 2:** Ion approaches of external binding pocket in K504 *in silico* mutants compared to *Os*HKT2;2-wt. The proportion of time which cations stay within 4Å of binding pocket forming residues. Time is expressed as a percentage during 100ns MD simulations. In addition, the number of approaches per ion species is indicated.

| (% of MD) | wt |  | K504R |  | K504Q |  | K504A |  | K504E |  |
| --- | --- | --- | --- | --- | --- | --- | --- | --- | --- | --- |
| Residue | Na<sup>+</sup> | K<sup>+</sup> | Na<sup>+</sup> | K<sup>+</sup> | Na<sup>+</sup> | K<sup>+</sup> | Na<sup>+</sup> | K<sup>+</sup> | Na<sup>+</sup> | K<sup>+</sup> |
| P71 | 1.9 | 0.4 | 0.5 | 1.4 | 6.0 | 4.4 | 1.9 | 0.3 | 3.2 | 1.0 |
| D75 | 1.4 | 0.2 | 0.1 | 0.6 | 10.0 | 26.8 | 8.4 | 6.9 | 27.4 | 27.7 |
| D501 | 1.8 | 0.5 | 2.5 | 3.2 | 12.9 | 13.8 | 9.6 | 7.3 | 16.0 | 11.6 |
| K504X | 2.3 | 0.2 | 0.2 | 0.4 | 8.0 | 21.2 | 0.02 | - | 24.0 | 38.7 |
| No. of approaches | 12 | 12 | 16 | 7 | 15 | 43 | 35 | 7 | 85 | 33 |

In *Os*HKT2;2-K504R, where the positive charge was kept but the side chain slightly increased, Na^+^, as well as K^+^, approached the binding pocket only shortly (see the short duration of stay and the low number of approaches in Table 2). Both cations were most of the time close to residue D501 (Na^+^: 2.5% and K^+^: 3.2% of simulation time), while in the wild type the duration of stay was more balanced between the four binding pocket residues. It is plausible that a more voluminous arginine residue at position 504 is preventing cations from entering into the binding pocket due to the repulsion of positive charges. Although, the surface area around the binding pocket is still negatively charged and able to attract cations, the positive charge at position 504 is in the case of the mutant bigger and further exposed to the extracellular space than in the wild-type channel, which may cause repulsion of cations (Figure 8E, J compared to A, F).

**Figure 8:**
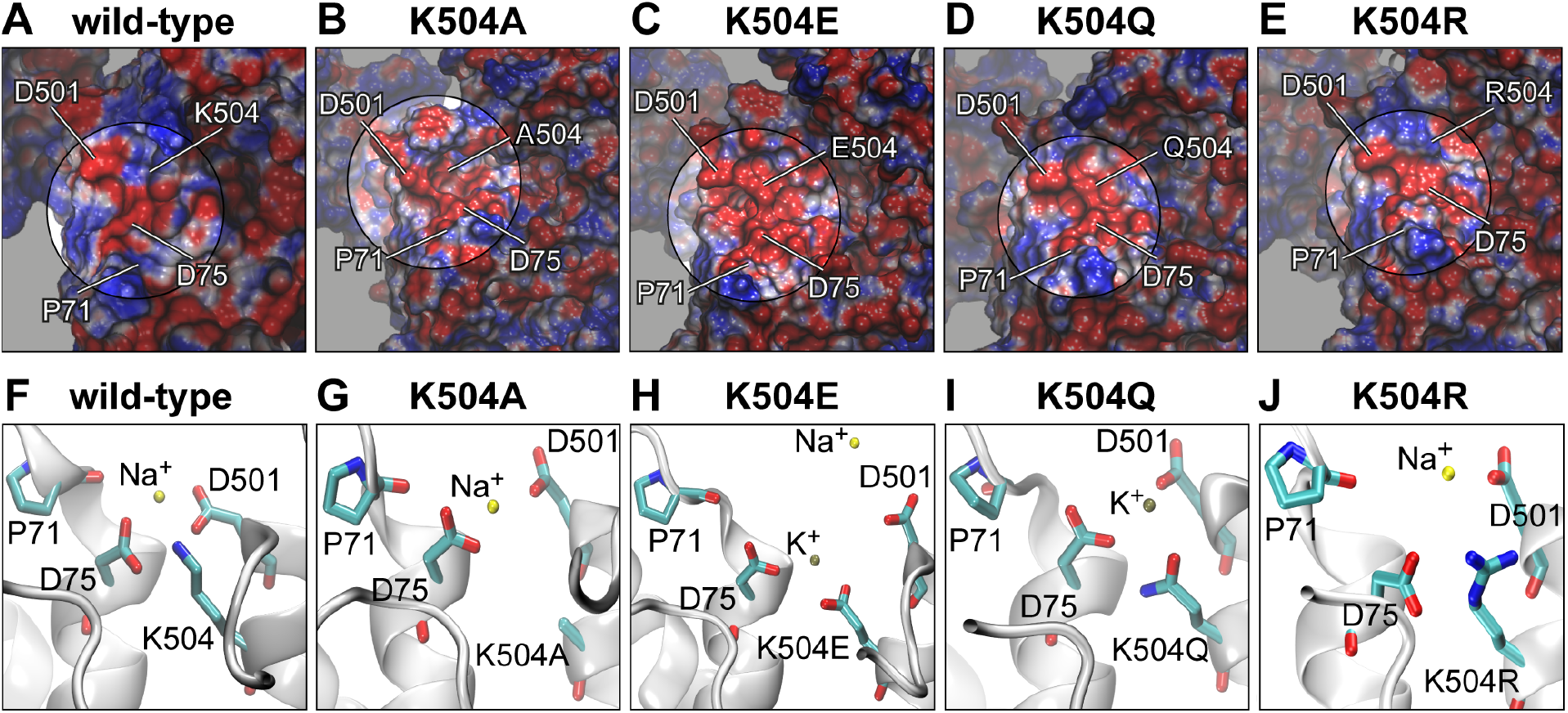
Binding pocket constitution of *in silico* mutants K504A, K504E, K504Q and K504R. (**A-E**) Electrostatic potential mapped to the protein surface. The cation binding pocket is highlighted and the position of P71, D75, D501 and K504 are indicated. (**F-J**) Representative illustration of the binding pocket constitution of all four K504 mutants and the wild type with ions approaching from the extracellular site or being inside the binding pocket.

In mutant K504Q, where lysine’s charge was neutralized, cations stayed at least five times longer in the binding pocket (12.9% and 26.8% for Na^+^ and K^+^, respectively) than in mutant K504R, although both mutants showed comparable ion transport abilities in electrophysiological experiments. In contrast to mutant K504R, cations entered into the binding pocket in mutant K504Q, which may account for the more extended stay in the pocket (compare Figure 8I, J). Additionally, K^+^ ions approached the binding pocket six times more frequently than in K504R, which may account at least in part for the longer duration of stay of K^+^ in the binding pocket. Compared to the wild type, cations stayed significantly longer close to the binding pocket. A similar increase (at least for Na^+^ ions) was observed in mutant K504A. The absence of the positive charge may allow cations to stay longer in the negatively charged binding pocket before continuing to the pore entrance of the channel.

In the non-functional mutants K504A and K504E, Na^+^ approached the binding pocket with high frequency (35 and 85 times) and stayed up to 9.6% (K504A) and 27.4% (K594E) of the simulation time. K^+^ ions stayed shorter time than Na^+^ ions in the binding pocket in mutant K504A (up to 7.3%) and longer in mutant K504E (up to 38.7%). The insertion of a negatively charged glutamate in mutant K504E generated a large negative area on the protein surface compared to *Os*HKT2;2-wt (Figure 7C compared to A). A negative electrostatic potential highly attracts cations, and more than one cation approached the ion binding pocket at a time (Figure 7G). The long duration of stay indicates that the ions get trapped in the binding pocket, which may account for the absence of ion conduction in *Os*HKT2;2-K504E.

In summary, *in silico* data provide insights into potential reasons for functional alterations and the non-functioning of K504 mutants that are in good agreement with experimental data.

## 4 Discussion

HKT channels have been described as crucial players in salt tolerance. They transport Na^+^ as uniporter (class I) or in symport with K^+^ ions (class II). Class II-type HKT channels function as Na^+^ uniporter in the absence of K^+^. However, they become Na^+^/K^+^ symporter in the presence of K^+^. This mutual effect of ion-species on the ion conduction has been observed for several HKT channels. Dependent on the channel member the influence on ion transport varies (Jabnoune et al., 2009). Our electrophysiological studies confirmed the stimulating effect that K^+^ ions exert on Na^+^ transport in *Os*HKT2;2. Interestingly, this effect was only seen at low K^+^ concentrations and decreased as the extracellular K^+^ concentration increased. This phenomenon has been previously reported for *Os*HKT2;2 (Oomen et al., 2012). From the physiological point of view, this is consistent behavior. During K^+^ starvation (low K^+^ concentrations), ion uptake increases and with it the uptake of Na^+^ ions. Physico-chemically similar Na^+^ ions may complement functions of K^+^ up to a certain degree and may enable the survival of the plant while K^+^ availability is low. With increasing concentration of K^+^, the ion conduction is reduced and with it the uptake of Na^+^ ions. In the presence of sufficient K^+^ ions, high Na^+^ influx is not desired since Na^+^ ions compete with K^+^ and may cause symptoms of K^+^ starvation. However, while the physiological response of HKT is well investigated, the ion-specific effect at the molecular level is still unknown.

To get deeper insights into the molecular basis behind the mutual ion-species effect, we modeled the structure of *Os*HKT2;2 and analyzed cation-protein contacts. Our computational studies show that four residues P71, D75, D501 and K504 form an extracellular cation binding pocket. Thereby, P71, D75 and D501 coordinate cations and the positively charged lysine (K504) holds the two negatively charged aspartates, D75 and D501, in place. Thus, a negative environment that attracts cations is formed.

We confirmed the relevance of the binding pocket for the ion conduction of *Os*HKT2;2 in functional electrophysiological studies. Out of the three residues involved in ion coordination, residue D501 is slightly more sensitive towards modifications. D75 and P71 are located nearby on one extracellular end of a transmembrane helix and contribute to the cation coordination with each one oxygen atom. D501 is placed on the opposite side on the extracellular end of another helix and coordinates with two oxygen atoms. Therefore, mutating D501 removes all coordinating atoms from this side of the binding pocket, while mutating D75 is only eliminating one of two coordinating atoms from the opposite side, which may account for the slightly stronger effect seen in D501 mutants. In either case, these mutations maintain the overall functionality of *Os*HKT2;2 and affect its kinetics only slightly. Considering this binding pocket as one of many regions that ions pass before entering the pore, it is plausible that mutations affect the channel kinetics instead of rendering the channel non-functional. The identified binding pocket would be involved in the attraction of cations and their forwarding towards the pore entrance, rather than in the tight and constant binding of ions. Therefore, modifying the coordinating oxygen atoms would first affect transport dynamics.

Lysine 504, on the other hand, is a crucial residue in the binding pocket. Inverting the charge from positive to negative by a glutamate substitution results in a non-conductive mutant. With a third negative charge in the binding pocket, the two aspartates repulse and drift apart, which leads to an expansion of the binding pocket. Additionally, the three negative charges form a large negative region on the protein surface that attracts cations and traps them in the binding pocket. If the identified binding pocket is understood as one stopover for ions on their way into the pore, the trapped ions may explain the inhibited ion conduction, since they might block the path for ions leading into the pore.

Overall, the positively charged lysine 504 may be the necessary impulse that cations need to be forwarded towards the pore. The positive charge and moreover its correct size and position are crucial for channel functionality. The sensitivity towards changes of one of the parameters size or charge indicates how fine-tuned the constitution of the binding pocket is. Removing the side chain by mutations to alanine renders *Os*HKT2;2 seemingly non-functional. Neutralizing the positive charge (K504Q) or increasing the size of the amino acid side chain with simultaneous maintenance of its charge (K504R), led to substantial changes in the conductance of *Os*HKT2;2. In the latter, the Na^+^ uniport is strongly reduced at high Na^+^ concentrations. However, at low K^+^ concentrations, these mutants still mediate large ion currents indicating that either the binding pocket still works correctly for K^+^ ions or that the positive effect of K^+^ ions has another origin. One possibility may be, e.g. the existence of another ion binding pocket that is more relevant for the K^+^ forwarding than the one described here. In fact, during simulations, we could identify additional regions that are approached by ions on their way into the pore. These regions are currently under investigation.

The importance of K504 has been shown earlier for the wheat TaHKT1 channel (Kato et al., 2007). In this study, the mutant K508Q, which corresponds to K504Q in our present study, showed reduced functionality in comparison to the wild-type channel in electrophysiological experiments and yeast complementation studies. The authors also proposed a potential salt bridge formation between K508 and D78, which corresponds to D75 in *Os*HKT2;2. Those results and the ones presented in our study point to the functional relevance of K504 in combination with D75. Our study additionally provides a biological and structural context by identifying these residues as part of a cation binding pocket.

Overall, using computational methods, we identified an extracellular cation binding pocket and validated its functional relevance experimentally. Although being about 20Å far from the pore entrance, the identified binding pocket is essential for proper ion conduction. Essential ion binding pockets on the extracellular protein surface provide an example of how different ion species may affect the channel conduction behavior even before entering the pore. We are still far from understanding how K^+^ ions influence the ion transport of *Os*HKT2;2 on the molecular level. This study, however, provides new evidence, which opens the discussion for new explanations for the mutual effect of ion species.

## 5 Acknowledgments

We thank Dr. Julian Schroeder (University of California, San Diego) for providing the clone of *Os*HKT2;2 wild-type. We also thank Alison Coluccio for technical assistance. This work was supported by grants from the Chilean Fondo Nacional de Desarrollo Científico y Tecnológico (http://www.conicyt.cl/fondecyt) to JR (No. 3150173), AV-J (No. 11170223), ID (No. 1150054) and WG (No. 1140624), and from Conicyt, programme PAI, Convocatoria nacional subvención a la instalación en la academia 2017 to JR (No. PAI77170035).

## 7 Author contributions

JR conceived and coordinated the study and wrote the paper. JR and AVJ designed, performed and analyzed the computational experiments. JR and MP designed, performed and analyzed the electrophysiological experiments. JR, ID and WG interpreted the data. All authors reviewed the results, revised the manuscript and approved its final version.

